# IL-23 receptor signaling licenses group 3-like innate lymphoid cells to restrict a live-attenuated oral Chlamydia vaccine in the gut

**DOI:** 10.1101/2023.09.11.557246

**Authors:** Ying He, Yihui Wang, Rongze He, Ahmed Mohamed Abdelsalam, Guangming Zhong

**Author notes:** Corresponding author: Guangming Zhong, Department of Microbiology, Immunology and Molecular Genetics University of Texas Health Science Center at San Antonio, 7703 Floyd Curl Drive, San Antonio, Texas 78229, USA.

## Abstract

An IFNγ-susceptible mutant of *Chlamydia muridarum* is attenuated in pathogenicity in the genital tract and recently licensed as an *intr*acellular *O*ral vaccine *v*ector or intrOv. Oral delivery of intrOv induces transmucosal protection in the genital tract but intrOv itself is cleared from the gut (without shedding any infectious particles externally) by IFNγ from group 3-like innate lymphoid cells (ILC3s). We further characterized the intrOv interactions with ILC3s in the current study since the interactions may impact both the safety and efficacy of intrOv as an oral Chlamydia vaccine. Intracolonic inoculation with intrOv induced IFNγ that in return inhibited intrOv. The intrOv-IFNγ interactions were dependent on RORγt, a signature transcriptional factor of ILC3s. Consistently, transfer of oral intrOv-induced ILC3s from RORγt-GFP reporter mice to IFNγ-deficient mice rescued the inhibition of intrOv. Thus, IFNγ produced by intrOv-induced ILC3s is likely responsible for inhibiting intrOv, which is further supported by the observation that oral intrOv did induce significant levels of IFNγ-producing LC3s (IFNγ^+^ILC3s). Interestingly, IL-23 receptor knockout (IL-23R^-/-^) mice no longer inhibited intrOv, which was accompanied with reduced colonic IFNγ. Transfer of oral intrOv-induced ILC3s rescued the IL-23R^-/-^ mice to inhibit intrOv, validating the dependence of ILC3s on IL-23R signaling for inhibiting intrOv. Clearly, intrOv induces intestinal IFNγ^+^ILC3s for its own inhibition in the gut, which is facilitated by IL-23R signaling. These findings have provided a mechanism for ensuring the safety of intrOv as an oral Chlamydia vaccine and a platform for investigating how oral intrOv induces transmucosal protection in the genital tract.

**Significance:** Despite the extensive efforts, no subunit vaccine is available for protecting humans against *C. trachomatis* infection and pathogenicity. Recent studies have led to a licensed live-attenuated oral vaccine (intrOv) that is both effective in inducing transmucosal protection in the genital tract and safe due to its susceptibility to IFNγ delivered by ILC3s. Characterization of the intrOv interactions with IFNγ-producing ILC3s in the current study has revealed a critical role of IL-23 receptor signaling in facilitating ILC3s to clear intrOv from the gut, which has provided a mechanism for ensuring the safety of intrOv and laid a foundation for further revealing the mechanisms by which oral intrOv induces transmucosal immunity in the genital tract since ILC3s can also function as antigen presenting cells.

## Introduction

Sexually transmitted infection with *Chlamydia trachomatis* may lead to pathological sequelae in the upper genital tract (1–4). Although effective antibiotics are available for treating chlamydial infection, the challenge is that *C. trachomatis* infection is often asymptomatic or lacks specific symptoms, thus significantly hindering the treatment efficacy. Obviously, vaccination offers a long-term solution for this challenge. However, despite the extensive efforts in more than half-century (5–23), there is still no licensed Chlamydia vaccine for humans (14). The failure of inactivated trachoma vaccine trials carried out after the isolation of ocular chlamydia (6, 7, 24–26) had motivated the search for a subunit *C. trachomatis* vaccine, for which hundreds of chlamydial antigens have been evaluated in preclinical models (13, 16, 27). One subunit vaccine construct was recently evaluated in a phase I trial(19) but its efficacy in humans is still to be evaluated (17). Even if this vaccine is ultimately approved for human use, there may still be a need to develop other versions of Chlamydia vaccines given the diverse human populations affected by Chlamydia. For example, both IPV (inactivated polio vaccine) and OPV (oral live polio vaccine) have contributed to the world-wide eradication of the polio pandemic and their combinational usage is continuing to maintain human herd immunity (28). Further, although mRNA vaccines have been successful in ending the COVID-19 pandemic in many countries, a live-attenuated mucosal vaccine may still be needed for blocking the spreading of SARS-CoV-2 and preventing the emerging of new variants (29). Thus, developing whole organism-based vaccines against Chlamydia is both timely and necessary. In 2015, the inactivated *C. trachomatis* organisms were modified to induce protective immunity in preclinical models (18), indicating that the whole chlamydial cell-based vaccine approach can be considered. However, producing a *C. trachomatis* serovar-based vaccine at large scales and simultaneously combating against all 8 genital *C. trachomatis* serovars may be challenging for this vaccine. Live attenuated *C. psittaci* (30) & *C. abortus* (31) vaccines are approved for protecting animals since the animal-adapted chlamydial organisms are much easier to produce in large scales. The yield of the mouse-adapted *C. muridarum* organisms is estimated at ∼100 folds higher than that of the human pathogen *C. trachomatis* under similar culture conditions, which makes large scale production of *C. muridarum* possible.

*C. muridarum* has been used for investigating chlamydial pathogenicity in the genital tract (32), which is also known to establish long-lasting colonization in the mouse gastrointestinal (GI) tract. The GI tract *C. muridarum* not only fails to cause pathologies (33–37) but also induces transmucosal immunity against subsequent infection in the genital tract (38) and airway (39). Antibiotic clearance of GI *C. muridarum* does not affect the protective immunity in the genital tract (38), indicating that the gut-primed immunity can be sustained in extra-gut tissues. These surprising findings made during our pathogenesis studies reignited our interest in developing *C. muridarum* into an attenuated live oral vaccine for protecting humans against *C. trachomatis*. This is because there are successful examples of using live attenuated vaccines to induce cross-species or heterotypic protection in humans such as cowpox virus as the vaccine against smallpox (40) and BCG of *M. bovis* against *M. tuberculosis* (41, 42). A meningococcal vaccine has recently been found to protect against gonorrhoea (43). Consistently, mice with prior exposure to *C. muridarum* developed heterotypic protection against subsequent infection with *C. trachomatis* (44), suggesting that the mouse-adapted *C. muridarum* may also be used to protect humans against *C. trachomatis*.

To improve the safety of a *C. muridarum*-based vaccine, various genital tract pathogenicity-attenuated *C. muridarum* clones have been identified (45–48). Although these mutants are no longer pathogenic in the genital tract (49, 50), they still retain the capacity to induce immunity against genital infection. One variant is designated as <u>intr</u>acellular <u>O</u>ral vaccine <u>v</u>ector (intrOv) that is now licensed as an oral Chlamydia vaccine (48). While IND-enabling studies for supporting clinical trials of intrOv are underway, the current study is to continue to investigate intrOv interactions with gut mucosal tissues, which may provide mechanistic information on how to improve both the safety and efficacy of intrOv. A unique property of intrOv is that a single oral inoculation can induce a robust transmucosal protection in the genital tract (48) but no live intrOv organisms can be recovered from the rectal swabs (51, 52). The latter makes intrOv a safe oral vaccine. The inhibition of intrOv in the gut was mapped to the large intestine colon, which is why intrOv was also defined as a colon colonization-deficient mutant (52, 53). Further studies revealed that intrOv was rescued to regain long-lasting colonization in the colon of IFNγ^-/-^ mice, which led to the designation of intrOv as an IFNγ-susceptible mutant (52). Using the inhibition of intracolonically inoculated intrOv as a model, recent studies have identified the cellular source of the colonic IFNγ responsible for inhibiting intrOv (51, 54, 55). It was found that the responsible IFNγ must be provided by innate cells since Rag1^-/-^ mice failed to rescue intrOv colonization but further depletion of IFNγ from Rag1^-/-^ mice did so. The most likely innate cells might be the group 3-like innate lymphoid cells (ILC3s) since deficiency in RORγt, the signature of transcriptional factor of ILC3s, also rescued intrOv colonization. However, many questions on whether or how intrOv induces IFNγ-producing ILC3s (IFNγ^+^ILC3s) and how IFNγ^+^ILC3s inhibit intrOv remain unknown.

ILC3s, expressing the transcriptional factor RORγt and cytokines IL-17 and IL-22, consist of multiple subpopulations with dynamic/diverse functions(56, 57). ILC3s can respond to diverse signals from microbes, food and host via various receptors, including Ffar2(58), Ah receptor (AhR; ref:(59), IL-1R(60), IL-18R(61) and IL-23R(62). ILC3s differentiate into different stages for producing IFNγ, becoming IFNγ^+^ILC3s or ex-ILC3s (63, 64). Particularly, IL-23 has been shown to be critical in activating ILC3s to secrete IL-22 and driving ILC3s to produce IFNγ (65–67). It is worth emphasizing that IL-23 is a multi-functional cytokine and IL-23R has been shown to drive apoptosis of Tregs in the gut (68). Since chlamydial infection is known to induce the expression of many cytokines including IL-23 (69–72), It will be interesting to evaluate whether IL-23R signaling plays a critical role in the intrOv induction of IFNγ^+^ILC3s.

In the current study, we further characterized the intrOv interactions with ILC3s in the gut. It was found that intracolonic inoculation with intrOv significantly induced IFNγ which was accompanied by reduced intrOv, both of which were dependent on RORγt, the signature transcriptional factor of ILC3s. Consistently, when oral intrOv-induced ILC3s from the intestinal lamina propria of RORγt-GFP reporter mice were adoptively transferred to the IFNγ-deficient recipient mice, the intracolonically inoculated intrOv was inhibited, indicative of IFNγ production by the intrOv-induced ILC3s. Oral delivery of intrOv was indeed found to induce significant levels of IFNγ-producing LC3s (IFNγ^+^ILC3s) in the intestinal lamina propria. To determine the role of IL-23 signaling in intrOv interactions with ILC3s, mice deficient in IL-23 receptor (IL-23R^-/-^) were evaluated using the intracolonic inoculation model. It was found that IL-23R^-/-^ no longer inhibited intrOv and also significantly reduced colonic IFNγ. However, the IL-23R^-/-^ mice were rescued to inhibit intrOv by adoptive transfer with oral intrOv-induced ILC3s, demonstrating the dependence of ILC3s on IL-23 signaling for inhibiting intrOv. The above observations have demonstrated that intrOv induces IFNγ^+^ILC3s for its own inhibition, which is facilitated by IL-23 receptor signaling. These findings have provided both a mechanism for ensuring the safety of the attenuated Chlamydia vaccine intrOv and a platform for further revealing the mechanisms of oral intrOv-induced transmucosal protection in the genital tract since ILC3s can also function as antigen-presenting cells (73, 74).

## Results

### 1. The oral vaccine intrOv induces colonic IFNγ and is inhibited by IFNγ in the colon

We have previously demonstrated that intrOv is deficient in colonizing mouse colon (52, 53) due to its susceptibility to IFNγ (52). ILC3s appeared to be both necessary and sufficient for inhibiting intrOv colonization in the mouse colon (51, 54). However, it remains unclear whether and how intrOv induces ILC3s to produce IFNγ for restricting its own colonization in the mouse colon. In the current study, we first monitored colonic IFNγ production over time following intracolonic inoculation with intrOv (Fig. 1). It was found that in wild type mice, both IFNγ mRNA and protein production peaked on day 7, which correlated with significant reduction in intrOv recovery, indicating that intrOv can induce colonic IFNγ which may in return inhibit intrOv replication. However, this induction and inhibition loop was reversed in mice deficient in RORγt (Fig. 2). It was found that homozygous mice with EGFP KI (*gfp* gene replacing Rorcγt, defined as RORγt-deficient mice or RORγt^-/-^) were significantly compromised in producing IFNγ with a level of <30 pg/ml of colon tissue homogenates on day 7 after intracolonic inoculation with intrOv. However, under the same experimental condition, the heterozygous mice (that still maintain a copy of the wild type Rorcγt) or the wild type C57 mice were able to produce >60 pg of IFNγ per ml of colon homogenate tissues. This >50% reduction in the level of colonic IFNγ reversely corelated well with a significant increase in the yields of intrOv in both the colon tissues and rectal swabs. More importantly, the colonic IFNγ reduction in RORγt^-/-^ mice as also translated into the rescue of the long-lasting colonization by intrOv in the colon as shown in Fig. 3. Although the intracolonically inoculated intrOv was cleared within 2 weeks in the RORgt-EGFP heterozygous mice or wild type C57 mice, intrOv maintained its shedding of live organisms in the rectal swabs for 4 weeks and significant levels of live intrOv organisms were detected in the large intestinal tissues when examined on day 28. These observations together have demonstrated that intrOv induction of IFNγ for its own inhibition in the colon is dependent on RORγt, a signature transcriptional factor of ILC3s.

**Fig. 1.**
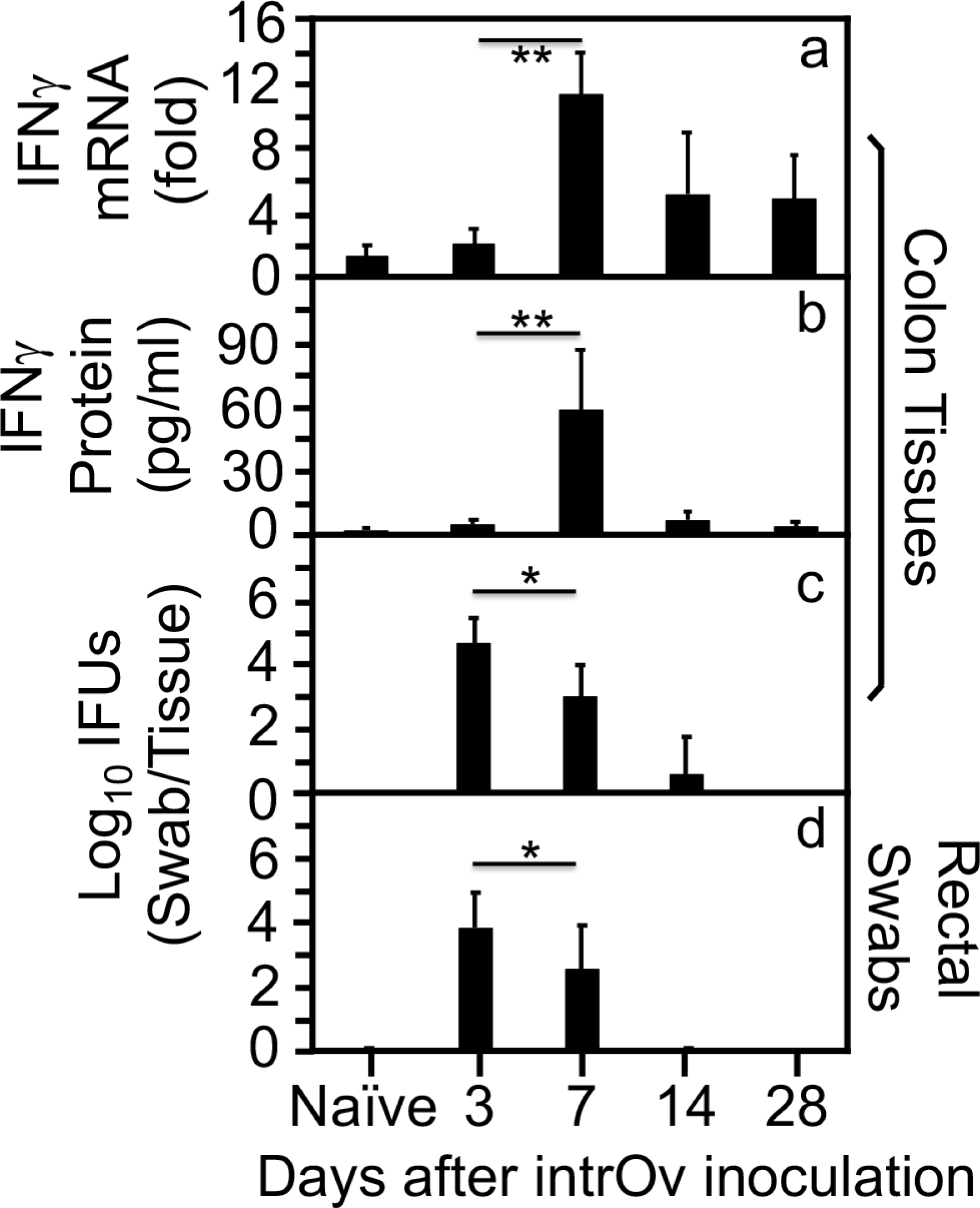
IFNγ production and intrOv yields following intracolonic inoculation with the attenuated and IFNγ-susceptible *C. muridarum* mutant intrOv. C57BL/6J mice without (naïve) or with intracolonic inoculation with intrOv at 1 x 10^7^ inclusion forming units (IFUs) were sacrificed for collecting colon tissues on days 3, 7, 14 & 28 as indicated along the X-axis for measuring IFNγ mRNA (panel a, change in fold over naïve mice, n=4 or 5), IFNγ protein (b, pg/ml, n=4 or 5) and live intrOv burden (c, Log_10_IFUs per colon tissue, n=4 to 5). Rectal swabs were also taken for monitoring intrOv shedding (d, Log_10_IFUs per swab, n=5). Note that colonic IFNγ mRNA and protein peaked on day 7, corelating with significant reduction in intrOv burden (*p<0.05, **p<0.01, 2-tailed Wilcoxon). The data were from two independent experiments.

**Fig. 2.**
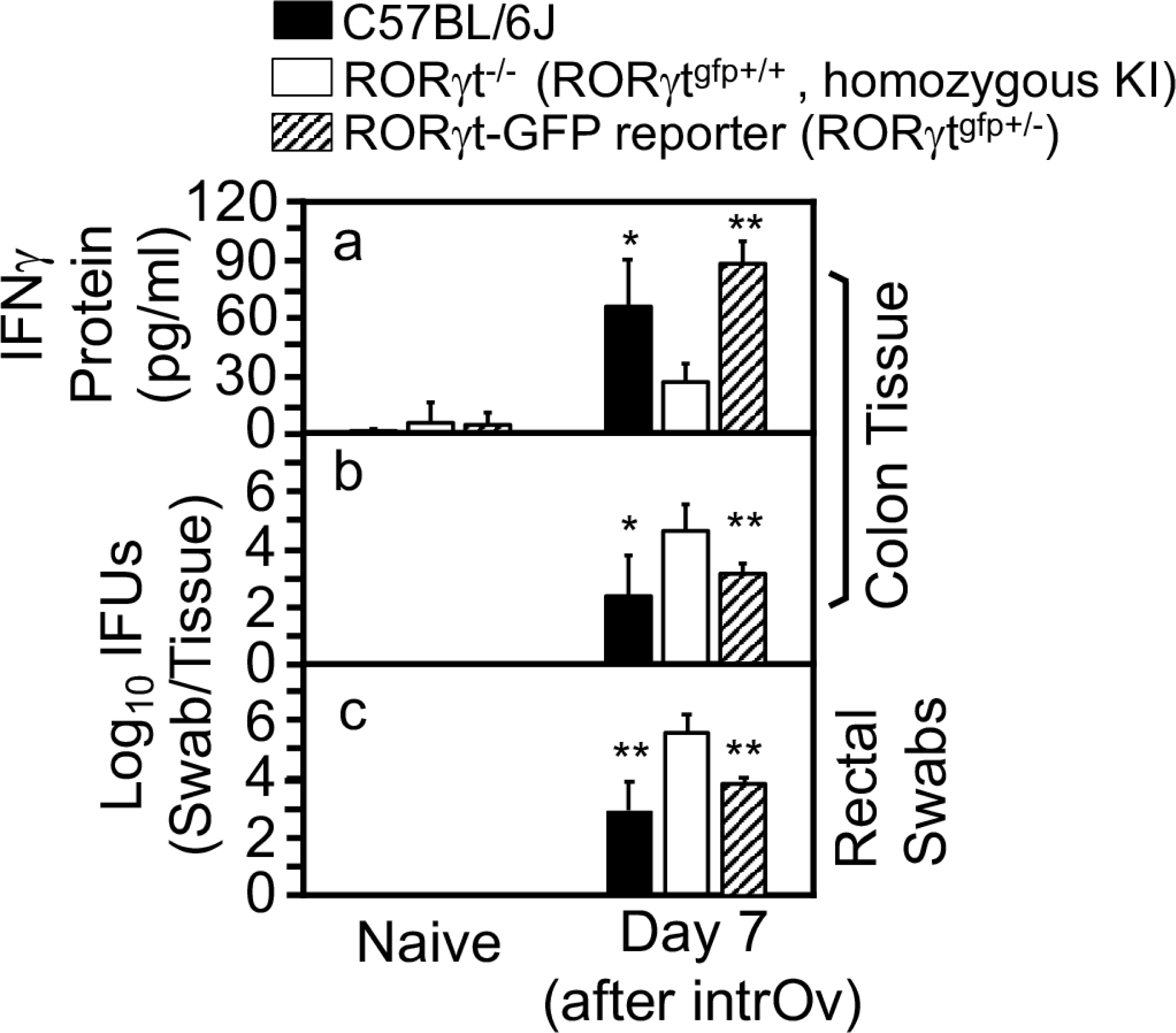
Comparison of IFNγ production and intrOv yields in colon tissue on day 7 after inoculation between mice with or without RORγt. C57BL/6J (solid bar), RORγt-eGFP knock-in (KI) homozygous mice (RORγt^gfp+/+^, which is deficient in RORγt or RORγt^-/-^, open bar) or heterozygous mice (RORγt^gfp+/-^ or RORγt-GFP reporter, hatched bar) were intracolonically inoculated without (naïve) or with intrOv at 1 x 10^7^ IFUs. On day 7, all mice were sacrificed for collecting colon tissues for measuring IFNγ protein (panel a, pg/ml, n=4) and intrOv burdens (b, Log_10_IFUs per colon tissue, n=4). Rectal swabs were also taken for monitoring intrOv shedding (c, Log_10_IFUs per swab, n=4). Note that colonic IFNγ was significantly induced in C57BL/6J and RORγt GFP reporter but not RORγt^-/-^ mice while intrOv yield significantly increased in RORγt^-/-^ mice. \**p*<0.05, \*\**P*<0.01, 2-tailed Wilcoxon. Data were from two independent experiments.

**Fig. 3.**
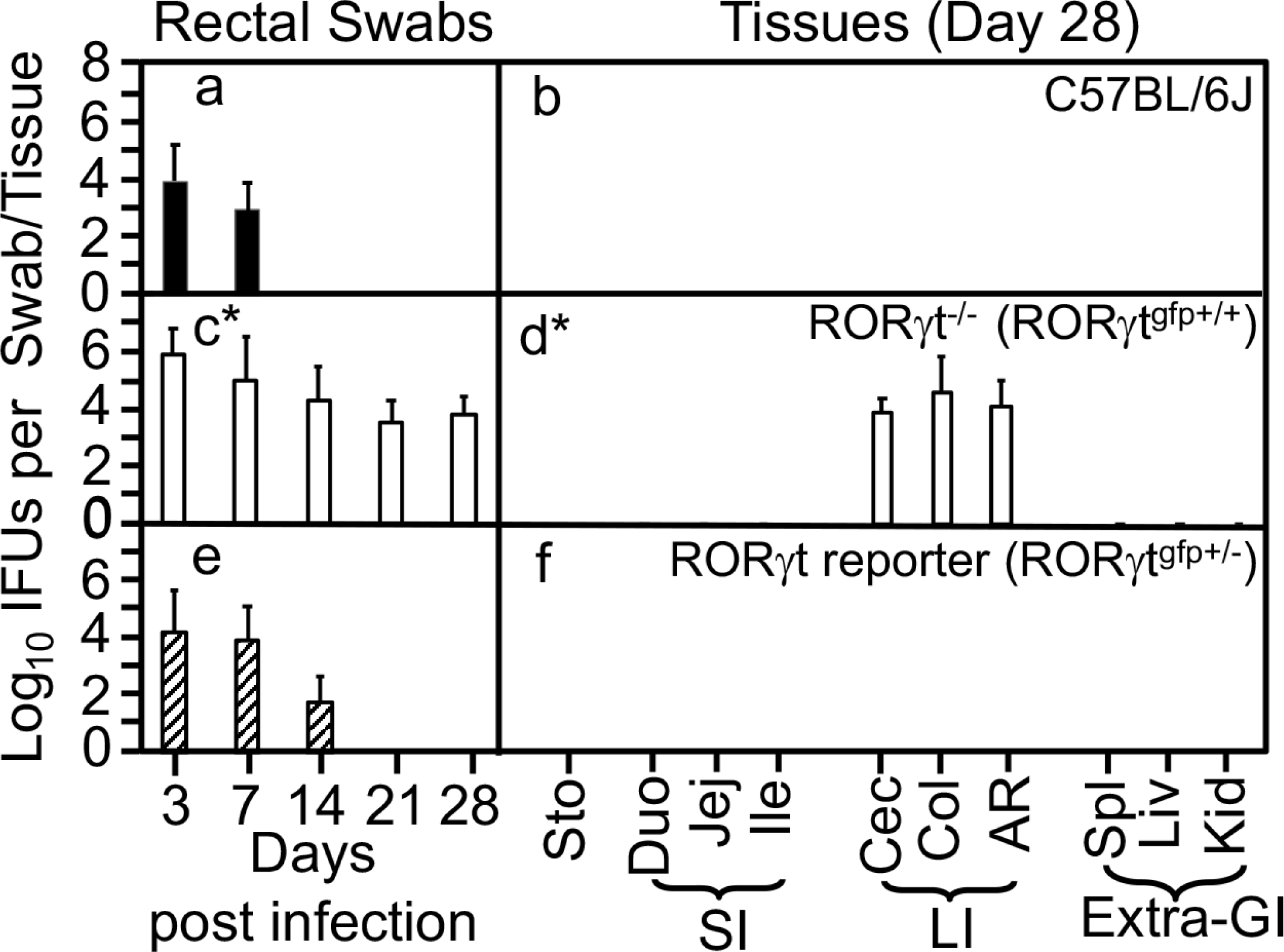
IntrOv yields recovered from mice with or without RORγt following intrOv inoculation. C57BL/6J (solid bar, panels a & b), RORγt-eGFP KI homozygous mice (RORγt^-/-^, open bar, c & d) or heterozygous RORγt GFP reporter mice (RORγt^gfp+/-^, hatched bar, e & f) were intracolonically inoculated with intrOv at 1 x 10^7^ IFUs. On days 3, 7 and weekly thereafter, rectal swabs were taken (a, c & e) or on day 28, mouse tissues were harvested (b, d & f) as indicated along the X-axis for monitoring intrOv yields. The mouse tissues include stomach (Sto), small intestinal tissues (SI) duodenum (Duo), jejunum (Jej), and Ilium (Ile), and large intestinal tissues (LI) cecum (Cec), colon (Col) and rectum (Rec), as well as extra-gastrointestinal tissues (Extra-GI) spleen (Spl), liver (Liv) and kidney (Kid) as indicated along the X-axis. The intrOv yields were expressed as Log_10_IFUs per swab or tissue, as shown along the Y-axis. Note that RORγt^-/-^ mice significantly increased intrOv yields comparing to either C57 or the reporter mice (n=5, Area-under-curves were used for comparison between groups using 2-tailed Wilcoxon, *p<0.05). Data were from two independent experiments.

### 2. The IFN**γ** responsible for inhibiting intrOv in the colon may be delivered by ILC3s

The RORγt-EGFP heterozygous mice significantly elevated colonic IFNγ and inhibited intrOv following intrOv inoculation while the homozygous mice (lacking RORγt) failed to do so (Figs 2 & 3), which has demonstrated that both the induction of colonic IFNγ by intrOv and the inhibition of intrOv by IFNγ are dependent on RORγt. Since the RORγt-dependent IFNγ is likely produced by ex-ILC3s, we further hypothesize that the heterozygous mice may be induced to develop IFNγ-producing ILC3s in the intestine by the oral vaccine intrOv. This hypothesis can be tested by isolating intrOv-induced ILC3s from the heterozygous mice and further evaluating their ability to inhibit intrOv colonization in recipient mice that are permissible to intrOv colonization. As shown in Fig. 4, significant number of live ILC3s can be isolated from the intestinal lamina propria of the heterozygous mice after oral inoculation with live intrOv for 7 days. After excluding dead cells and lineage-positive cells, ∼7% of the remaining live cells are positive for GFP. These are the authentic live ILC3s since they are positive for RORγt, which can be used as donor cells for subsequent adoptive transfer experiments. As shown in Fig.5, we used the sorter-purified live ILC3s as donor cells for transferring to IFNγ-deficient mice (IFNγ^-/-^). We chose IFNγ^-/-^ mice as recipient mice because we have previously shown that these mice allow intrOv to establish long-lasting colonization in the colon following an intracolonic inoculation (51, 52). It was found that twice transfer each with 40,000 ILC3s was sufficient for preventing intrOv from establishing long-lasting colonization in the colon of IFNγ^-/-^ mice while transfer with non-ILC3s failed to do so. It is reasonable to assume that the ILC3s donor cells must produce IFNγ for compensating the loss of IFNγ in the IFNγ^-/-^ mice.

**Fig. 4.**
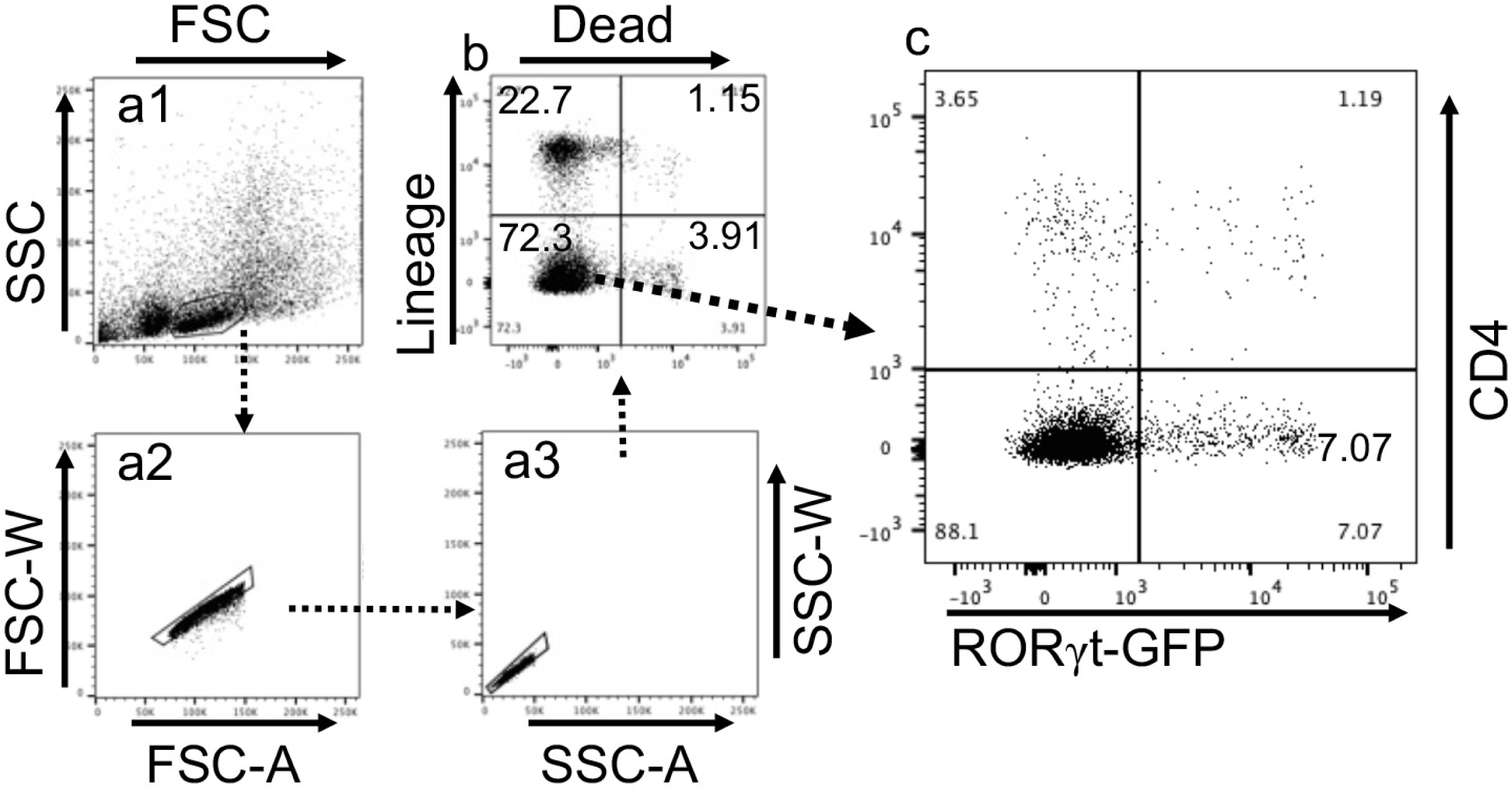
Optimizing sorter-purification of ILC3s from RORγt-GFP reporter mice orally primed with the oral vaccine intrOv. RORγt-GFP KI heterozygous mice (RORγt-GFP reporter mice) orally inoculated with 1 x 10^7^ IFUs of live intrOv for 7 days were sacrificed for harvesting intestinal lamina propria lymphoid cells (LPL). After ficoll density gradient centrifugation and immunostaining with surface markers, single cells were gated based on cell’s physical properties (a1-a3). Then, non-viable and lin+(positive for CD3, Ly-6G/Ly-6C, CD11b, CD45R/B220, & TER-119) cells were excluded (b). Finally, live GFP+ and CD4-negative cells were collected as ILC3s (c) for subsequent adoptive transfer experiments.

**Fig. 5.**
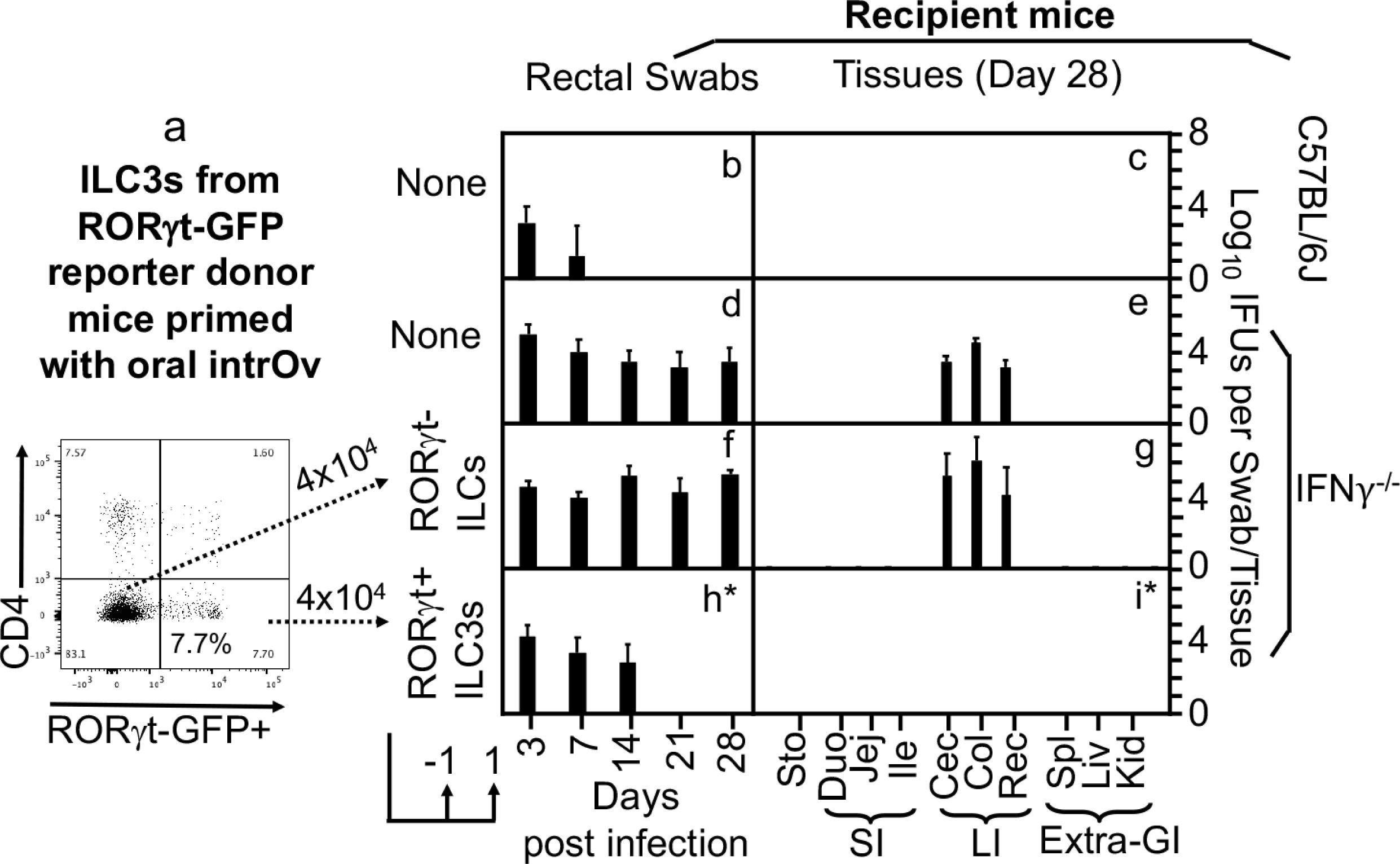
Adoptive transfer with RORγt GFP+ ILC3s vs. GFP-non-ILC3s as donor cells to restore inhibition of intrOv colonization in IFNγ-deficient recipient mice. RORγt GFP+ ILC3s and GFP-non-ILC3s (panel a) were sorter-purified from donor mice as described in figure 4 legend above as donor cells for adoptively transferring to IFNγ-deficient recipient mice. The transfer was carried out twice with 4 x 10^4^ cells for each and on one day before (-1) and one day after (+1) the intracolonic inoculation of recipient mice as indicated at the bottom. A dose of 1 x 10^7^ IFUs of live intrOv was used to intracolonically inoculate C57BL/6J mice (b & c, n=4) or IFNγ-/- mice without (d & e, n=4) or with transfer of RORγt GFP-non-ILC3s (f & g, n=3) or RORγt GFP+ ILC3s (h & I, n=3). On days 3, 7 and weekly thereafter after inoculation, rectal swabs were taken (b, d, f & h) or on day 28, mouse tissues were harvested (c, e, g & i) as indicated along the X-axis for monitoring live chlamydial organisms. The mouse tissues include stomach (Sto), small intestinal tissues (SI) duodenum (Duo), jejunum (Jej), and Ilium (Ile), and large intestinal tissues (LI) cecum (Cec), colon (Col) and rectum (Rec), as well as extra-gastrointestinal tissues (Extra-GI) spleen (Spl), liver (Liv) and kidney (Kid) as indicated along the X-axis. The yields of intrOv expressed as Log_10_IFUs per swab or tissue were as shown along the Y-axis on the right. Note that transfer of ILC3s but not non-ILC3s conferred resistance to intrOv colonization in IFNγ-/- mice. *p<0.05, Wilcoxon rank-sum (Area-under-curves between IFNγ-/- transferred with RORγt GFP-non-ILC3s versus RORγt GFP+ ILC3s). Data were from two independent experiments.

### 3. Oral intrOv is able to induce IFNγ^+^ILC3s in intestinal lamina propria

By taking advantage of the RORγt-GFP reporter mice for sorter-purification of live ILC3s, we have demonstrated that adoptive transfer of oral intrOv-induced ILC3s from the reporter mice to IFNγ^-/-^ recipient mice can rescue the IFNγ^-/-^ mice to inhibit intrOv, suggesting that the oral intrOv-induced ILC3s must be able to secrete IFNγ. However, direct evidence for intrOv to induce IFNγ-producing ILC3s or IFNγ^+^ILC3s remains lack. Here, we will use Rag1^-/-^ mice for providing the direct evidence. As shown in Fig. 6, we have optimized a gating scheme for selectively enriching ILC3s from the Rag1^-/-^ mouse intestinal lamina propria lymphoid cells. After excluding dead cells and lineage+ cells, we have validated the utility of the CD45 vs. CD90 plot for identifying ILC3s-enriched cell population, which are the CD45^int^CD90^hi^ cluster cells (54, 75). Indeed, ∼95% of CD45^int^CD90^hi^ cluster cells are RORγt-positive ILC3s. We then used the same gating strategy to compare the yields of IFNγ^+^ILC3s between Rag1^-/-^ mice with or without oral intrOv (Fig. 7). It was found that although most CD45^int^CD90^hi^ cluster cells are RORγt-positive ILC3s regardless of the oral intrOv treatment, oral intrOv treatment significantly promoted RORγt-positive ILC3s to express IFNγ (becoming IFNγ^+^ILC3s): >10% of RORγt-positive ILC3s from oral intrOv-treated Rag1^-/-^ mice expressed IFNγ while <3% of RORγt-positive ILC3s from oral SPG buffer-treated Rag1^-/-^ mice did so. Thus, although oral intrOv doesn’t significantly alter the proportion of ILC3s in mouse intestinal LPL, oral intrOv can significantly elevate the % of IFNγ^+^ILC3s.

**Fig. 6.**
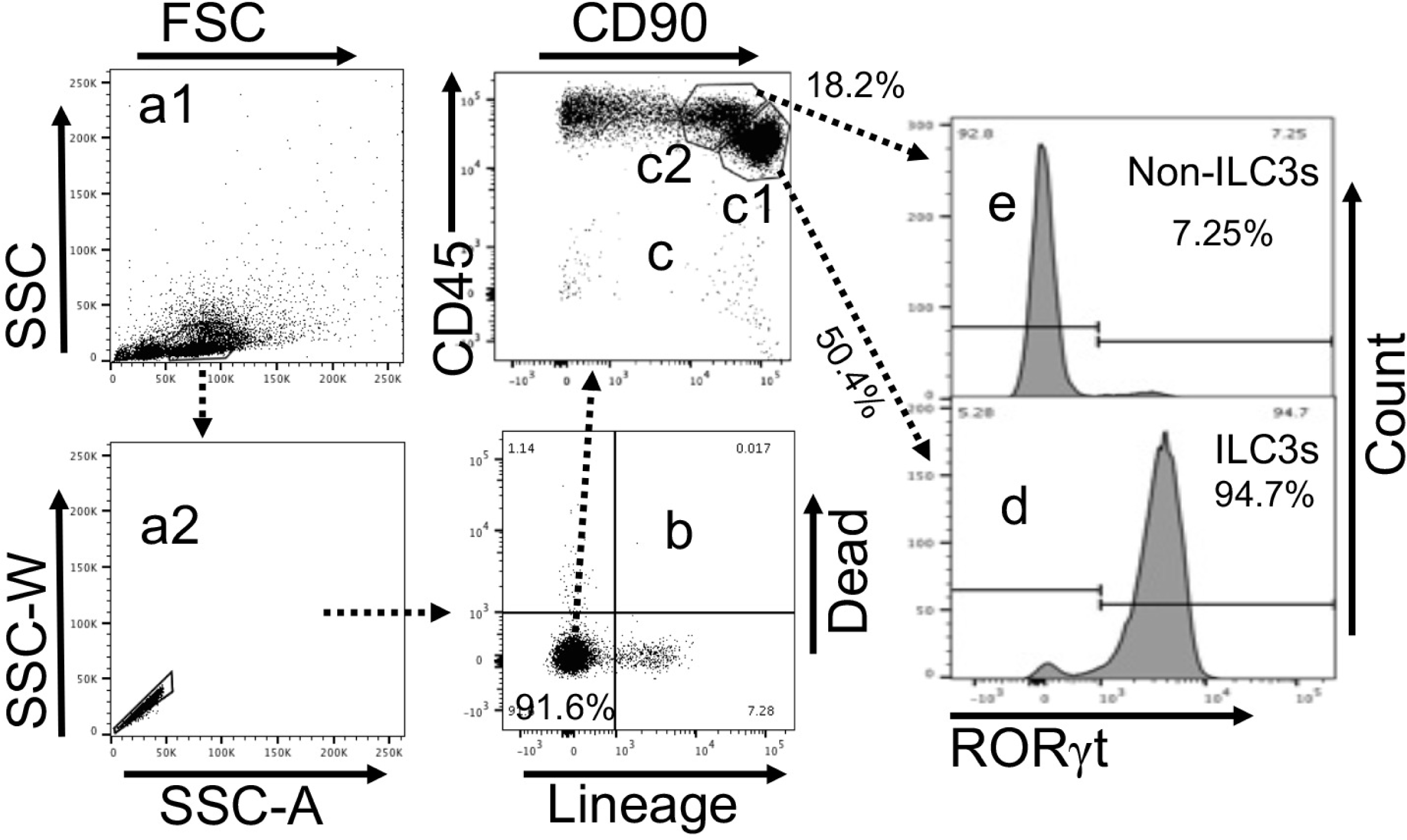
Optimizing sorter-purification of ILC3s from Rag1^-/-^ mice. Rag1-/- mice orally incubated with 1 x 10^7^ IFUs of live intrOv for 7 days were sacrificed for harvesting intestinal lamina propria lymphoid cells (LPL). After Ficoll density gradient centrifugation and immunostaining with antibodies against surface markers and the intracellular RORγt, single cells were gated based on cell’s physical properties (a1-a2). Then, dead cells and lin+(positive for CD3, Ly-6G/Ly-6C, CD11b, CD45R/B220, & TER-119) cells were excluded (b). Further, the remaining live lineage-negative cells were plotted for the levels of CD45 & CD90 (c) where the CD45 intermediate and CD90 high (CD45^int^CD90^hi^) cluster cells (c1) were collected as ILC3s while the CD45 high and CD90 high (CD45^hi^CD90^hi^) cluster cells (c2) as non-ILC3s. Consistently, ∼95% of CD45^int^CD90^hi^ cells are positive for RORγt, a signature transcriptional factor of ILC3s while only ∼7% of CD45^hi^CD90^hi^ cells are RORγt-positive. The sorter-purified live CD45^int^CD90^hi^ and CD45^hi^CD90^hi^ cells from the same donor mice were used as donors for subsequent adoptive transfer experiments.

**Fig. 7.**
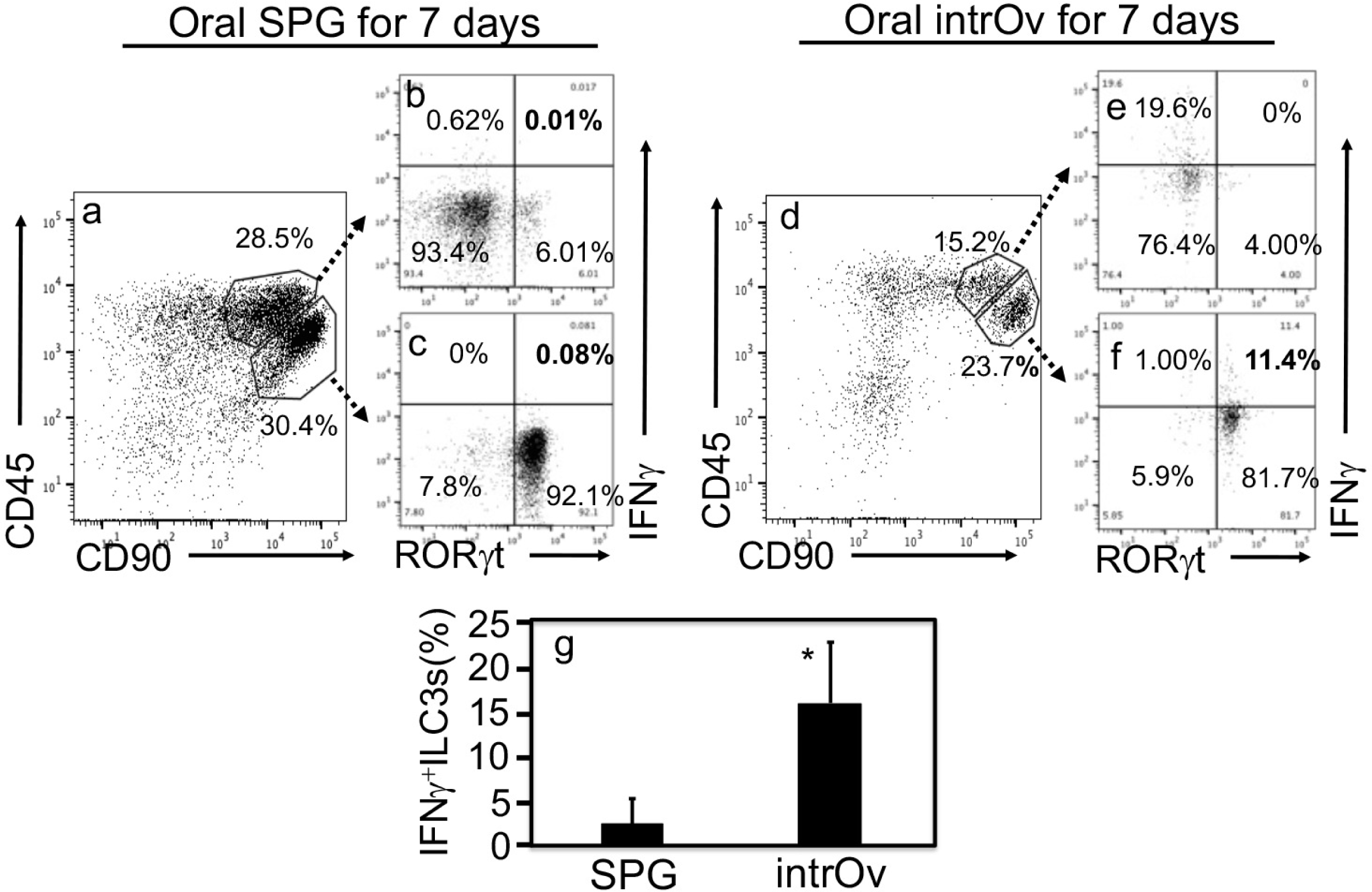
Oral intrOv induction of IFNγ-producing ILC3s. Rag1-/- mice orally incubated with 1 x 10^7^ IFUs of live intrOv (panels d-f) or SPG buffer alone (a-c) for 7 days were sacrificed for harvesting intestinal lamina propria lymphoid (LPL) cells. After Ficoll density gradient centrifugation and stimulation with mitogen PMA for 3 hours in the presence of a Golgi stopper, LPL cells were immunostained with antibodies against surface markers and intracellular IFNγ and RORγt. After gating for single cells based on cells’ physical properties and excluding dead and lineage-positive cells as described in Fig. 8 legend above, the CD45 vs. CD90 plot was used to identify the ILC3s-enriched CD45^int^CD90^hi^ cluster cells and no-ILC3s CD45^hi^CD90^hi^ cluster cells (panels a & d). Finally, both cluster cells were gated for intracellular RORγt and IFNγ (b, c, e & f). Note that oral intrOv significantly increased IFNγ-producing ILC3s (g, n=5, 2-tail Wilcoxon, *p<0.05).

### 4. IL-23R signaling is necessary for both inhibiting intrOv and inducing IFNγ^+^ILC3s

Having demonstrated the oral intrOv induction of IFNγ^+^ILC3s, the next question is to address the how intrOv induces IFNγ^+^ILC3s. Since IL-23 signaling is known to promote the differentiation of ILC3s to produce IFNγ (76, 77) and chlamydial infection has been shown to induce IL-23 (69, 70), we evaluated the role of IL-23 signaling in intrOv interactions with colonic ILC3s by using the homozygous IL-23R-EGFP KI mice (deficient in IL-23R or IL-23R^-/-^). As shown in Fig.8, IL-23R^-/-^ mice and wild type C57 mice were compared for susceptibility to intrOv colonization following an intracolonic inoculation. In the wild type mice, intracolonically inoculated intrOv was rapidly cleared. By the way, we previously found that no live organisms can be recovered from rectal swabs of wild type mice following an oral inoculation with intrOv while live organisms can be transiently detected from rectal swabs following an intracolonic inoculation (51–53). Thus, we have been using oral inoculation with intrOv to induce IFNγ^+^ILC3s and intracolonic inoculation to evaluate the inhibition of intrOv. In IL-23R^-/-^ mice, the intracolonically inoculated intrOv established long-lasting colonization in the large intestine. Thus, IL-23R^-/-^ mice allowed intrOv to achieve robust growth and establish long-lasting colonization in the large intestine while the wild type mice restricted intrOv colonization, demonstrating that IL-23 signaling is essential for restricting the colonization of intrOv. Further, the increased susceptibility of IL-23R^-/-^ mice to intrOv colonization corelated well with the failure of IL-23R^-/-^ mice to significantly elevate IFNγ production in response to the induction by intrOv (Fig. 9). Thus, IL-23R signaling is essential for both intrOv induction of IFNγ and inhibition of intrOv by IFNγ.

**Fig. 8.**
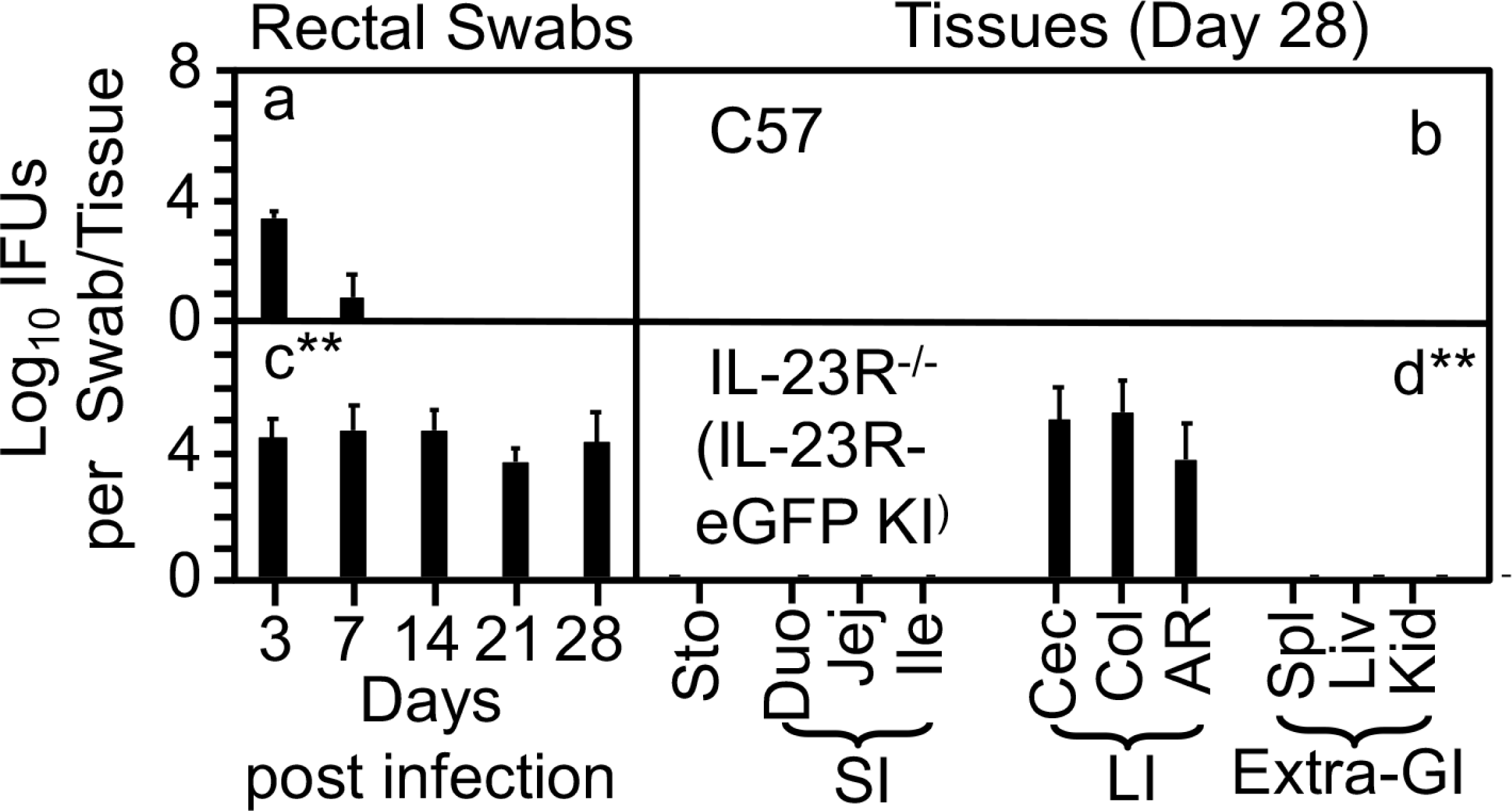
IntrOv burdens in mice with or without deficiency in IL-23 receptor. C57BL/6J (panels a & b, n=4) or homozygous IL-23 receptor-eGFP knock-in mice (deficient in IL-23R or IL-23R^-/-^, c & d, n=4) were intracolonically inoculated with intrOv at 1 x 10^7^ IFUs. On days 3, 7 and weekly thereafter after inoculation, rectal swabs were taken (a & c), and on day 28, all mice were sacrificed and tissues were harvested (b & c) as indicated along the X-axis for monitoring live intrOv burdens. The individual mouse tissues were described in figure 5 legend above. The intrOv burdens expressed as Log_10_IFUs per swab or tissue were shown (along the Y-axis). Note that IL-23R^-/-^ mice significantly increased intrOv colonization and allowed intrOv to establish long-lasting colonization in the large intestine (**p<0.01, 2-tail Wilcoxon comparison of Area-under-curves between IL-23R^-/-^ and C57BL/6J groups). Data were from two independent experiments.

**Fig.9.**
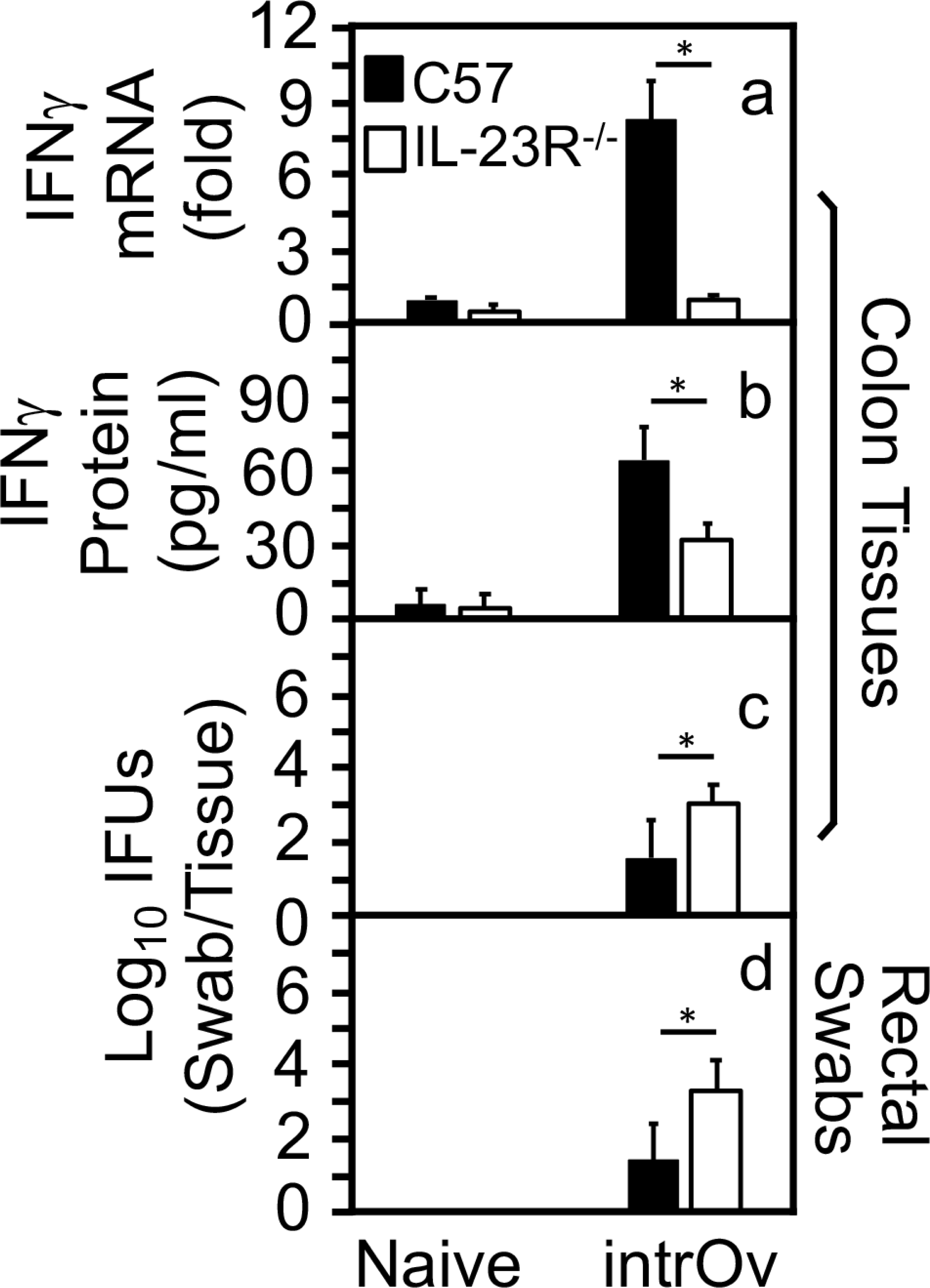
IFNγ production and intrOv yields following intrOv intracolonic inoculation between mice with or without deficiency in IL-23 receptor. C57BL/6J (C57, solid bar) or IL-23 receptor-deficient mice (IL-23R^-/-^, open bar) were intracolonically inoculated without (naïve) or with intrOv at 1 x 10^7^ IFUs. On day 7 after inoculation, mice were sacrificed for collecting colon tissues for measuring IFNγ mRNA (panel a, change in fold, n=3 for each naïve and 4 for intrOv-inoculated groups respectively), IFNγ protein (b, pg/ml, n=4) and live intrOv burden (c, Log_10_IFUs per colon tissue, n=4). Rectal swabs were also taken for monitoring intrOv burden (d, Log_10_IFUs per swab, n=4). Note that colonic IFNγ was significantly induced in C57BL/6J mice but not IL-23R^-/-^ mice, reversely corelating with intrOv burdens (\**p* < 0.05, Wilcoxon rank-sum between C57 & IL-23R^-/-^ groups). Data were from two independent experiments.

### 5. Oral intrOv-induced ILC3s are sufficient for rescuing IL-23R^-/-^ to inhibit intrOv

Having demonstrated the capacity of oral intrOv induction of IFNγ^+^ILC3s and the necessity of IL-23R signaling for both intrOv induction of IFNγ and intrOv inhibition by IFNγ, we next evaluated whether oral intrOv-induced ILC3s could rescue IL-23R^-/-^ mice to inhibit intrOv (Fig. 10). The CD45^int^CD90^hi^ cluster cells sorted from the intestinal lamina propria lymphoid cells isolated from Rag1^-/-^ mice 7 days after oral intrOv priming were used as IL-23R-competent donor ILC3s for adoptive transfer. The similarly sorted non-ILC3 cells from the same donor mice were used as control donor cells. The IL-23R^-/-^ mice were used as recipient mice since we wanted to test whether the IL-23R-deficiency can be compensated by IL-23R-competent donor ILC3s. The non-ILC3s sorted from the same donor mice were used as control donor cells. It was found that the IL-23R^-/-^ recipient mice that received the non-ILC3s remained susceptible to intrOv colonization since significant levels of live intrOv were continuously recovered from rectal swabs along the time course and the large intestinal tissues by day 28 when the tissues were harvested. However, IL-23R^-/-^ mice that were transferred with ILC3s became resistant to intrOv colonization. The IFUs from the rectal swabs decreased significantly on both days 7 and 14 and completely cleared by day 21. No live intrOv was detectable in any mouse tissues by day 28. Thus, the oral intrOv-induced ILC3s from Rag1^-/-^ mice are sufficient for inhibiting intrOv colonization in IL-23R^-/-^ mice.

**Fig. 10.**
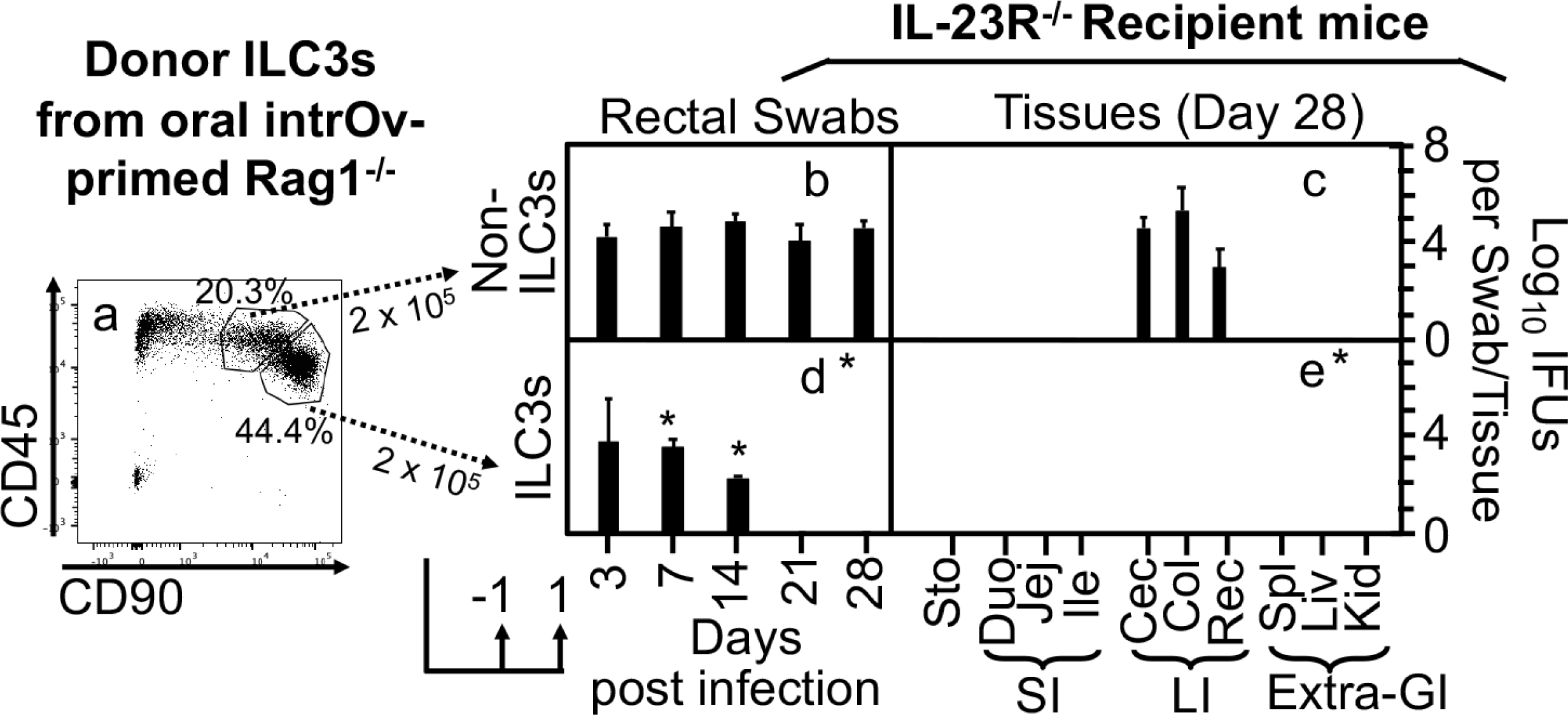
Adoptive transfer of intrOv-induced ILC3s from Rag1^-/-^ donor mice to IL-23R^-/-^ recipient mice to rescue the inhibition of intrOv. Sorter-purified ILC3s (panels d & e) and non-ILC3s (b & c) from intrOv-primed donor Rag1-/- mice (a) were adoptively transferred to IL-23R-/- recipient mice. The transfer was carried out twice with 2 x 10^5^ cells for each and on one day before (-1) and one day after (+1) the recipient mice were intracolonically inoculated with intrOv, as indicated at the bottom. On days 3, 7 and weekly thereafter after inoculation, rectal swabs were taken (b & d) or on day 28, mouse tissues were harvested (c & e) as indicated along the X-axis for monitoring live intrOv burdens. The yields of live intrOv expressed as log_10_IFUs per swab or tissue were shown along the Y-axis. Note that transfer of ILC3s but not non-ILC3s conferred resistance to intrOv colonization in IL-23R^-/-^ mice. *p<0.05, Wilcoxon (Area-under-curves between recipient mice receiving non-ILC3s vs. ILC3s, n=5 from two independent experiments).

## Discussion

IntrOv is both attenuated in genital tract pathogenicity (49, 50) and deficient in colonizing the GI tract (52, 53) due to its susceptibility to IFNγ (51, 52), both of which ensure the safety of intrOv as an attenuated vaccine. The IFNγ responsible for clearing intrOv from the GI tract has been identified as innate IFNγ likely delivered by ILC3s (51, 54, 55). In the current study, the intrOv interactions with ILC3s in the gut were further characterized. First, the time courses of colonic IFNγ production and live intrOv recoveries from the same mouse gut tissues were inversely correlated following intracolonic inoculation with intrOv, suggesting that intrOv may induce IFNγ for its own inhibition. Second, both the induction of IFNγ and the inhibition of intrOv were dependent on RORγt, a signature transcriptional factor of ILC3s, suggesting that ILC3s may be required for producing IFNγ responsible for inhibiting intrOv. Third, oral intrOv did induce RORγt^+^ILC3s in the intestinal lamina propria and adoptive transfer of the intrOv-induced ILC3s as donor cells rescued IFNγ^-/-^ recipient mice to inhibit intrOv, indicating that intrOv may induce ILC3s to produce IFNγ for inhibiting intrOv. Consistently, oral intrOv did induce significant levels of IFNγ-producing ILC3s (IFNγ^+^ILC3s) in the intestinal lamina propria. Fourth, IL-23R^-/-^ mice no longer inhibited intrOv colonization, which was accompanied by reduced levels of IFNγ, demonstrating that IL-23R signaling is necessary for both the inhibition of intrOv and induction of IFNγ. Finally, IL-23R^-/-^ mice were rescued to inhibit intrOv by adoptive transfer of oral intrOv-induced ILC3s, validating the dependance of ILC3s on IL-23R signaling for inhibiting intrOv. Thus, the current study has demonstrated a critical role of IL-23R signaling in facilitating ILC3s inhibition of intrOv in the gut.

ILC3s are known to respond to diverse signals to differentiate into different stages for producing IFNγ, becoming IFNγ^+^ILC3s or ex-ILC3s. For example, *Y. enterocolitica* (63) and *C. difficile* (64) may induce ILC3s to secrete IFNγ. The mucosal ex-ILC3s may contribute to both protection and pathology during infections with microbes such as *Salmonella* (78) or *Toxoplasma* (79). However, the precise mechanism by which each infection condition induces IFNγ^+^ILC3s remains unclear. Nevertheless, receptors such as Ffar2 (58), AhR (59), and IL-1R (60), IL-18R (61) or IL-23R (62) have been shown to promote ILC3 differentiation. *Chlamydia* is an obligate intracellular bacterium that has to replicate inside epithelial cells. In the current study, IL-23R was found to be essential for chlamydial induction of IFNγ^+^ILC3s, which is consistent with previous findings that chlamydial infection can induce IL-23 (70–72, 80) and IL-23 is potent in driving ILC3s to produce IFNγ (65). Since a sustained exposure of ILC3s to IL-23 was found to be sufficient for activating STAT4 to induce type 1 polarization (66, 67), it is likely that the intracellular intrOv may continuously stimulate the infected or adjacent host cells to release IL-23 that can then maintain the conversion of ILC3s to IFNγ^+^ILC3s.

Although intrOv is inhibited and cleared by IFNγ^+^ILC3s in the colon, its parental wild type strain *C. muridarum* is known to maintain long-lasting colonization there (34–36, 81). Following an inoculation with *C. muridarum* at any mucosal tissue site, *C. muridarum* can spread systemically (35, 82) via the blood circulation (81, 83–85). However, the bacteremia is cleared within 2 weeks and *C. muridarum* can only persist in the large intestine for long periods of time (34, 36). Apparently, *C. muridarum* may both induce and evade the intestinal IFNγ^+^ILC3s responses, which is consistent with the property of other commensal bacteria in the gut. Bacteria that can maintain long-lasting colonization in the gut become part of the gut microbiome. Microbiota species are known to confer colonization resistance to pathogenic infections but the same resistance is evaded by the microbiota species themselves (86–88). A major colonization resistance mechanism is by maintaining a base level of IFNγ (87). Since the wild type *C. muridarum* does not cause any significant pathology in the GI tract despite its long-lasting colonization in the gut (38), *C. muridarum* may also be a commensal species in the mouse GI tract, contributing to the maintenance of base level of IFNγ by inducing IFNγ^+^ILC3s via activating IL-23R signaling. Consistently, C. muridarum is known to induce IL-23 in mice (69, 70). Thus, the *C. muridarum*-IL-23-IFNγ^+^ILC3s pathway may represent one of the mechanisms by which *C. muridarum* has adapted to mouse GI tract tissues.

The discovery of the *C. muridarum*-IL-23-IFNγ^+^ILC3s pathway was made possible with the availability of the IFNγ-susceptible mutant intrOv (49–52). There are also other IFNγ-susceptible mutants of *C. muridarum* (89, 90), some of which are also attenuated in the genital tract pathogenicity and can induce transmucoal immunity in the genital tract (46). It will be interesting to compare these IFNγ-susceptible mutants to determine whether they are all inhibited by the IL-23R signaling-dependent IFNγ^+^ILC3s in the colon. More importantly, the IL-23R signaling-dependent IFNγ^+^ILC3s pathway in the gut may also contribute to the oral chlamydia-induced transmucosal immunity in the genital tract since ILC3s can function as antigen presenting cells(74, 91–93) and ILC3s-produced IFNγ can drive Th1 phenotype(94). It will be important in testing the hypothesis that oral intrOv may use the IL-23R signaling pathway to induce IFNγ^+^ILC3s in the gut for priming adaptive immunity via presenting chlamydial protective epitopes and promoting Th1 development in the genital tract.

## Materials and Methods

### 1. *Chlamydia* organisms

The *Chlamydia muridarum* mutant clone G28.51.1 was used in the current study (49, 50). Due to its lack of both pathogenicity in the genital tract (49, 50) and colonization in the colon (52, 53), G28.51.1 has been proposed as an intracellular oral vaccine vector or intrOv (32). IntrOv contains two mutations: a mutation in gene *tc0237* leading to the 117^th^ codon change from glutamine (Q) to glutamic acid (E) or TC0237Q117E and a mutation in *tc0668,* resulting in a stop codon at 216 that originally codes for glycine (G) or TC0668G216*. These mutations have been demonstrated to be responsible for both the attenuated pathogenicity in the mouse genital tract (50, 53) and reduced colonization in the gastrointestinal tract (51–54). The intrOv organisms were grown up in HeLa cells (human cervical carcinoma epithelial cells; ATCC# CCL-2) for density gradient purification into elementary bodies (EBs) and the purified EBs were stored in aliquots @ -80°C until use.

### 2. Mouse infection

The mouse experiments were carried out following the recommendations in the Guide for the Care and Use of Laboratory Animals endorsed by the National Institutes of Health. The protocol was approved by the Committee on the Ethics of Laboratory Animal Experiments of the University of Texas Health Science Center at San Antonio.

The following 5-7 weeks old male or female mice were from Jackson Laboratories, Inc., Bar Harbor, ME: Wild type mice (C57BL/6J, stock No: 000664), Rorc(γt)-EGFP heterozygote or homozygote (B6.129P2(Cg)-Rorctm2Litt/J, 007572) or mice deficient in recombination activating gene 1 (B6.129S7-Rag1tm1Mom/J, 002216, or Rag1^-/-^), IFNγ (B6.129S7-Ifngtm1Ts/J, 002287, or IFNγ^-/-^) as well as the IL-23 receptor-eGFP knock-in (KI) homozygote mice (*Il23r^tm1Kuch^*/J, 035863) as IL-23 receptor deficient mice or IL-23R^-/-^. The Rorc(γt)-EGFP heterozygote mice were used as RORγt-GFP reporter mice while the homozygous littermates as RORγt knockout/deficient mice or RORγt^-/-^. All mice were inoculated without or with intrOv EBs at a dose of 1 × 10^7^ inclusion forming units (IFUs) per mouse via intracolonic or intragastric inoculation as described previously (48, 54, 95). Briefly, EBs were diluted in 50μl (for intracolonic) or 100μl (for oral) of SPG (220 mM sucrose, 12.5 mM phosphate, 4 mM l-glutamic acid, pH 7.5) buffer and delivered to the stomach or colon using a straight ball-tipped needle designed for mouse oral gavage (N-PK 020; Braintree Scientific, Inc., Braintree, MA). After inoculation, mice were monitored for live organism shedding in rectal swabs or sacrificed for titrating live organisms in designated organs/tissues.

### 3. Titrating live chlamydial organisms from mouse swabs and tissues

To monitor live chlamydial organisms shedding from GI tracts, rectal swabs were collected in 0.5 ml of SPG buffer and vortexed with glass beads to release infectious EBs for quantitation. For titrating live organisms from mouse tissues, organs/tissues, as indicated in individual experiments, were collected (on designated days as specified in individual experiments) to 0.5 ml (for each segment of the genital tract tissue) or 2ml (for each remaining tissues/organ) of SPG buffer, followed by homogenization and brief sonication. Live organisms in the supernatants were titrated on HeLa cells in duplicate. The total number of IFUs/swab or tissue was converted into log_10_ for calculating the group mean and standard deviation. Please note that the detection limits of the titration method are 10 IFUs per swab and 40 IFUs per tissue sample. A total of 50μl from each sample without (neat) or with serial dilutions were used to inoculate monolayer HeLa cells. The entire culture well was counted for chlamydial inclusions when the inclusion density was low, allowing the detection of a single IFU per 50μl sample.

### 4. Immunofluorescence assay

The immunofluorescence assay for visualizing and counting chlamydial inclusions in the Chlamydia-infected HeLa culture was described previously (96). Briefly, infected HeLa cells grown on coverslips were fixed with paraformaldehyde (Sigma, St. Louis, MO 63178) and permeabilized with saponin (Sigma). The monolayers were labeled with a rabbit anti-chlamydial antibody (raised by immunization with C. muridarum EBs) and a goat anti-rabbit IgG conjugated with Cy2 (green, Jackson ImmunoResearch Laboratories, Inc) to visualize chlamydial inclusions while a Hoechst dye (blue; Sigma) was used for labeling nuclear DNA. The labeled cell samples were viewed under an Olympus IX-80 fluorescence microscope equipped with multiple filter sets (Olympus, Melville, NY).

### 5. Quantitative reverse transcriptase polymerase chain reaction (qRT-PCR) for detecting mouse *ifnγ* expression

For some experiments, mouse colon tissues were used for measuring IFNγ mRNA and protein (see ELISA below) in addition to quantitating live chlamydial organisms (see above). After sacrificing mouse, the colon was excised and cut into 5 equal portions after rinsing off fecal matter from the longitudinally opened colon. 1/4 of each portion were collected for each colon for RNA extraction while the remaining colon tissues for making tissue homogenates in SPG as described above. To extract total RNA, the colon tissue pieces were further minced before adding to a Lysing Matrix D tube that contains beads (Cat#: 6913050, MP Biomedicals, Santa Ana, CA). After quick mix by inverting the tube several times, immediately froze the sample tube in liquid. The frozen tubes can be processed right away or stored at -80°C for processing at a future time. Total RNA were isolated using TRIzol reagent (Cat#: 93289, Sigma) according to the manufacturer’s instruction. 1 ml TRIzol was added into each frozen Lysing Matrix D tube to pulverize the colon tissue with a homogenizer (Mini-Beadbeater, Biospectra, Stroudsburg, PA) by homogenizing 45 seconds twice. Then, 200 μL chloroform was added into each tube followed by vigorous vibration for 15-30 seconds; After incubating at room temperature for 5 min, the tubes were centrifuged at 12,000 g for 15 min at 4°C to separate the aqueous phase from the organic phase. Carefully transfer the top aqueous phase (about 300-400μl) into a new RNase-free 1.5ml tube followed by adding 550 μl isopropanol (by the way, all solutions and supplies used for qRT-PCR should be RNase-free). After a gentle mix and incubation at room temperature for 5min, the tubes were centrifuged at 12,000 g for 10 min at 4°C. A pellet at the base of each tube might be visible. After discarding the supernatant, 1 ml 75% ethanol in DEPC-treated water (Cat# AM9906, Ambion, Inc., Austin, TX) was used to rinse the tube (centrifugation at 7,500 g for 5 min at 4°C prior to pouring off the ethanol). The remaining pellet in each tube was air-dried inside a hood (5-10min) and resuspended in 20ul DEPC water for quantitating total RNA using a spectrophotometer. First-strand cDNA was synthesized from 200 ng of total RNA in a 20 μl reaction using the iScript™ cDNA Synthesis Kit (Cat#: 1708891, Bio-Rad). A SYBR green real-time PCR kit (Cat# 001752A, Bio-Rad) and a CFX96 Real-time Detection System (Bio-Rad) were used to run the qPCR reaction. The 2^-ΔΔ*CT*^ method was used for data analysis (97). The target gene is mouse *ifnγ* while the reference gene is *gapdh* from the same sample. The following primers were synthesized by Integrated DNA Technologies, Inc. (San Diego, CA) and used in the current study: IFNγ forward primer (5′-TCAAGTGGCATAGATGTGGAAGAA-3’) and reverse primer (5′-TGGCTCTGCAGGATTTTCATG-3’) and GAPDH forward primer (5’-ACCACAGTCCATGCCATCAC-3’) and reverse primer (5’-TCCACCACCCTGTTGCTGTA-3’).

### 6. Measurement of cytokines in mouse tissues using ELISA

The ELISA kit for measuring mouse IFNγ from colon tissues was purchased from Thermo Fisher Scientific (Cat# 88-7314-88, Waltham, MA). The colon tissues homogenized in SPG buffer as described above were added with a protease inhibitor cocktail (cat#78430, 100x stock, Thermo Fisher Scientific) at the final concentration of 2X. The neat homogenates were 2-fold serially diluted with PBS containing 2X inhibitor cocktail and the diluted samples were applied to 96-well plates precoated with a capture antibody. IFNγ binding was detected with a detection antibody plus detection reagent. The absorbance at 450 nm was detected with a Synergy H4 microplate reader (BioTek, Winooski, VT), and the results were expressed as pg/ml based on the standard curve obtained from the plate.

### 7. Adoptive transfer

Innate lymphoid cells (ILC3s) were prepared from either RORγt-GFP reporter or Rag1^-/-^ mice as donor cells for adoptive transfer experiments. The transfer was carried out twice, each with designated number of donor cells, as indicated in individual experiments. One transfer was done via retro-orbital injection on the day before and the other on the day after intracolonic infection of the recipient mice with intrOv. Following the intracolonic infection, live intrOv organisms were monitored in both rectal swabs and tissues.

The donor ILCs were prepared as described previously (54, 75). Briefly, after Rag1^-/-^ donor mice were orally inoculated with 1 x 10^7^ IFUs of intrOv for 7 days before the intestinal tissues were collected for lymphoid cell isolation from lamina propria. After the intraepithelial lymphoid cells were first removed, the remaining tissue pieces were used for isolating lamina propria lymphoid cells by mincing the tissues into 1-2 mm pieces. The minced tissue mixture was digested with both DNase I and Liberase followed by shaking to release lymphoid cells into the supernatants. The process can be repeated several times. After filtering and washing, the lymphoid cells-containing solutions were collected for Percoll purification. The lymphoid cell-containing interphase was harvested as lamina propria lymphoid cells or LPLs for flow cytometry sorting after excluding dead and lineage positive cells (gating for live lineage negative or lin^-^ cells). Both the ILC3s-enriched subset (CD45^int^CD90^hi^) and non-ILC3s subset (CD45^hi^CD90^hi^) were sorted as donor cells. Typically, ∼1 x 10^5^ LP-ILC3s were obtained from a single donor mouse. For some experiments, the donor ILC3s were harvested from RORγt-EGFP heterozygote mice. In this case, GFP was used to sort for RORγt-expressing cells. To elevate RORγt-GFP cells in mouse intestinal lamina propria, 1 x 10^7^ IFUs of intrOv were orally inoculated for 7 days before the intestinal tissues were collected for lymphoid cell isolation from lamina propria. Similarly, after excluding dead and lineage positive cells, GFP+ cells were sorted as donor cells.

### 8. Flow cytometry analysis and sorting

The following antibodies were used for labeling surface markers: rat anti-mouse CD16/32 (block the nonspecific binding to Fc receptors, clone:2.4G2, cat#:BE0307, Bio X Cell, West Lebanon, NH), rat anti-mouse Lineage cocktail (conjugated with Pacific blue, cat#133310, Biolegend, Inc, San Diego, CA) consisting of rat anti-mouse CD3 (clone: 17A2), rat anti-mouse Ly-6G/Ly-6C (clone: RB6-8C5), rat anti-mouse CD11b (clone: M1/70), rat anti-mouse B220 (clone: RA3-6B2) and rat anti-mouse Ter-119 (clone: Ter-119), rat anti-mouse CD45.2 (conjugated with APC, clone#104, cat#17-0454-82, Thermo Fisher Scientific, Inc, Waltham, MA) and rat anti-mouse CD90.2 (conjugated with FITC, clone#30-H12, cat# 105305, Biolegend, Inc). Dead cells were excluded by staining with Viability-Dye (conjugated with eFluor 506, cat#65-0866-14, Thermo Fisher Scientific). The antibody staining was permitted for 30min at 4°C.

FACS Aria II instrument (BD Biosciences) was used to sort the desired cell population as donor cells for adoptive transfer experiments. After excluding dead and lineage positive cells (CD3+Ly6C+CD11b+B220+Ter119+), ILC3-enriched subset (CD45intCD90hi) and non-ILC3 subset (CD45hiCD90hi) were sorted from Rag1-deficient mouse LP-ILCs into ILC3-enriched and non-ILC3 subsets. GFP+ cells were sorted from RORγt-GFP reporter mice.

### 9. Statistics

The numbers of live organisms in IFUs at individual data points or over a time course were compared using the Wilcoxon rank-sum test. Area-under-the-curve or AUC was used for comparing the time course or clusters of tissue sample data. When multiple groups were included in a given experiment, ANOVA was first used to determine whether there was an overall significant difference among all groups. Only when p<0.05 (ANOVA), were the differences between every two groups further analyzed using Wilcoxon.

